# Untangling the ecological signal in the dental morphology in the bat superfamily Noctilionoidea

**DOI:** 10.1101/2021.07.21.453269

**Authors:** Camilo López-Aguirre, Suzanne J Hand, Nancy B Simmons, Mary T Silcox

## Abstract

Diet has been linked to the diversification of the bat superfamily Noctilionoidea, a group that underwent an impressive ecological adaptive radiation within Mammalia. For decades, studies have explored morphological adaptations and diversity of noctilionoid bats to reveal traits associated with their ecological diversity. Surprisingly, despite such interest and recent application of novel techniques, ecomorphological studies have failed to fully resolve the link between diet and a critical component of the feeding apparatus: dental morphology. Using multivariate dental topographic analysis and phylogenetic comparative methods, we examined the phylogenetic, biological and ecological signal in the dental morphology of noctilionoid bats. Analysing the lower first molars of 110 species, we explored relationships between diet and dental morphology, accounting for three different dimensions of diet (guild, composition and breadth). Phylogenetic and size-dependent structuring of the dental topography data shows it does not correlate only to diet, highlighting the need to account for multiple sources of variation. Frugivorous noctilionoids have sharper molars than other previously reported frugivorous mammals. Nectarivorous noctilionoids showed reduced lower molar crown height and steepness, whereas animalivorous species had larger molars. Dietary composition suggested that the intensity of exploitation of a resource is also linked to different dimensions of dental morphology. Increasing carnivory positively correlated with MA, explaining the highest proportion of its variation, and increasing frugivory explained the highest proportion of variation in all other variables. Dietary breadth showed generalist species have sharper, more topographically-complex molars, whereas specialist herbivores and specialist animalivores fell at opposite ends in the range of tooth steepness and crown height. Together, the results suggest that adaptations affecting different attributes of dental morphology likely facilitated the dietary diversity and specialisation found in Noctilionoidea.

## Introduction

The bat superfamily Noctilionoidea (order Chiroptera) is one of the most ecologically diverse groups of modern mammals, having the broadest range of dietary adaptations within Chiroptera (Rojas et al., 2016). Representing ∼17% of modern chiropteran species diversity, noctilionoid bats encompass the entire dietary range found in the more than 1400 species of bats (Rojas et al., 2016; Simmons and Cirranello, 2021). Feeding ecology (diet and foraging behaviours) has been repeatedly linked to the taxonomic and morphological diversification of this group (Dumont et al., 2012; López-Aguirre et al., 2021a; Monteiro & Nogueira, 2011; Rojas et al., 2018; Rossoni et al., 2019). Stemming from an ancestral insectivorous diet, noctilionoid bats have adapted to feed on blood, seeds, terrestrial vertebrates, fish, fruit, leaves, nectar and pollen (Rojas et al., 2011). Bat dietary adaptations unique to Noctilionoidea include sanguivory (i.e. feeding on blood) in the subfamily Desmodontinae, and granivory (i.e. feeding on seeds) and folivory (i.e. feeding on leaves) in species of the subfamily Stenodermatinae (Duque-Márquez et al., 2019; Nogueira & Peracchi, 2003; Villalobos-Chaves et al., 2016). Levels of omnivory across a herbivory-to-carnivory gradient have also been described in Noctilionoidea and linked to macroevolutionary trajectories during its diversification (Rojas et al., 2018). Most of noctilionoid dietary diversity is concentrated in the Neotropical family Phyllostomidae, with the exception of piscivory which is exclusive to the family Noctilionidae, being common in one species (*Noctilio leporinus*) and perhaps occasionally in another *N. albiventris* (Hood & Pitocchelli, 1983; Brooke, 1994). Species of the families Furipteridae, Mormoopidae, Myzopodidae and Thyropteridae are broadly categorised as insectivores (Arbour et al., 2019; Rojas et al., 2018), whereas species of the family Mystacinidae exhibit a broad omnivorous diet apparently facilitated by terrestrial locomotion (Arkins et al., 1999). There are multiple instances of convergent evolution of some dietary habits within Noctilionoidea (e.g. carnivory, frugivory and nectarivory), highlighting the high ecological adaptability of this group (Datzmann et al., 2010; Santana & Cheung, 2016).

There are multiple dimensions to the dietary diversity of noctilionoid bats, complicating the study of its ecomorphology. Comparisons across dietary guilds have been the main tool to quantify morphological differences, reflecting broad similarities across species in the same guild (e.g. Arbour et al., 2019; Dumont et al., 2012; Monteiro & Nogueira, 2011; Santana & Cheung, 2016). Classifying species into dietary guilds can oversimplify an individual species’ diet (Rojas et al., 2012, 2018), potentially underestimating the variety of food items that species feed on (herein dietary breadth). For example, *Vampyrum spectrum*, commonly classified as a carnivore, also feeds on insects, fruit and nectar (Rojas et al., 2018; Santana & Cheung, 2016). Additionally, the extent to which species exploit a given resource represents a different dimension of the diet of species (herein diet composition), as it reflects levels of niche partitioning and specialisation across species along a specialisation gradient (Hall et al., *in press*; Rojas et al., 2018). For example, nectarivory is used to different degrees in phyllostomid species that are specialist nectarivores (e.g. *Musonycteris harrisoni*), omnivore herbivores (e.g. *Carollia perspicillata*) and omnivore animalivores (e.g. *Lophostoma silvicolum*), exemplifying different dietary niches (Hall et al., *in press*; Rojas et al., 2018). Articulating the multidimensionality of dietary adaptations within a single comparative framework can better elucidate its ecological and evolutionary drivers (DeCasien et al., 2017; Hall et al., *in press*; Makedonska et al., 2012).

The functional link between the morphological and dietary diversity of extant noctilionoids has been an active area of research for decades (Arbour et al., 2019; Dumont, 2007; Dumont et al., 2012; Freeman, 1981, 1988; Monteiro & Nogueira, 2011; Nogueira et al., 2009; Santana et al., 2011, 2012; Santana & Cheung, 2016). The main area of study with respect to the relationship between diet and form in noctilionoids has been the ecomorphology of the skull, with a strong correspondence being found between aspects of cranial anatomy and elements of the diet (Arbour et al., 2019; Freeman, 1981; Rossoni et al., 2017, 2019; Santana et al., 2012). Different morphological adaptations have been linked to the exploitation of specific resources: increased body size with carnivory (Freeman, 1988; Santana & Cheung, 2016), elongated rostrum with nectarivory (Arbour et al., 2019; Freeman, 1995), sharpened incisors and reduced post-canine dentition with sanguivory (Arbour et al., 2019; Freeman, 1988), and u-shaped mandible and shorter rostrum with durophagy in frugivory (Arbour et al., 2019; Freeman, 1988; Santana et al., 2012). Moreover, phenotypic specialisations to particular feeding ecologies (e.g. carnivory and nectarivory) have been found to converge across Noctilionoidea (Arbour et al., 2019; Freeman, 1995; Morales et al., 2019; Santana & Cheung, 2016). As such, foraging strategies represent a significant driver of phenotypic variation in bats, with feeding ecology correlating with both cranial and postcranial morphology (Gaudioso et al., 2020; López-Aguirre et al., 2021a; Morales et al., 2019). An emerging body of research has also recently begun to focus on molecular and morphological adaptations in sensory systems associated with adaptation to particular dietary niches (Arbour et al., 2021; Hall et al., *in press*; Leiser-Miller & Santana, 2020).

Despite steady interest in the ecomorphology of diet in bats, dental adaptations and variability in teeth across extant bats remain poorly known and understudied. Freeman (1984, 1988) summarized broad patterns of dental adaptations to diet across bats but primarily considered tooth proportions. Dental traits have often been used to attempt to reconstruct dietary habits of fossil bats (e.g. Hand, 1985; Hand et al., 2018; Simmons et al., 2016) but mostly in a qualitative rather than quantitative fashion. Other studies have looked at teeth and variation in more detail, but often with either very small sample sizes and/or in highly taxonomically restricted groups. For example, Dumont (1995) evaluated the relationship between enamel thickness and diet in six species of bats as part of a broader study; Santana et al. (2011) explored differences in teeth crown occlusal complexity between dietary guilds; Ghazali and Dzheverin (2013) evaluated correlations between dental form and diet in 9 species of *Myotis*; and a recent study examined dental variation in species of flying foxes and its correlation to diet and phylogenetic relationships (Zuercher et al., 2021). In this context, the goal of the current study was to evaluate dental variation in a much more ecologically diverse clade of bats using larger sample sizes and accounting for three different dimensions of diet (guild, composition, and breadth). To accomplish this, we used methods that serve to quantify multiple aspects of dental variation.

Dental Topographic Analysis (DTA) comprises an increasingly widely used set of metrics that aim to quantify diet-based dental adaptations (Berthaume et al., 2020; Evans et al., 2007; Ledogar et al., 2013; Selig et al., 2021; Ungar et al., 2018). Originally based on geographic information system analysis (e.g., Ungar & M’Kirera, 2003; Evans et al., 2007), DTA has revolutionised the understanding of how the dentition adapts to the biomechanical properties of foodstuff (Bunn et al., 2011; Ledogar et al., 2013; Melstrom & Irmis, 2019; Ungar et al., 2018). A robust set of DTA metrics has been developed in recent decades, allowing quantification of varying aspects of a complex adaptive landscape (Berthaume et al., 2019a, 2020; Boyer, 2008; Cooke, 2011; Evans et al., 2007; Pineda-Munoz et al., 2017; Winchester et al., 2014). In contrast to geometric morphometrics (another widespread set of morphometric tools), DTA is homology-free, facilitating comparison among a wide range of taxa and structures (Berthaume et al., 2020; Melstrom & Irmis, 2019; Pineda-Munoz et al., 2017). Dental topographic analysis was developed and has been used primarily to study mammal dentitions, most regularly applied to study Primates (e.g., Berthaume et al., 2020; Boyer, 2008; Bunn et al., 2011; Evans et al., 2007; Pineda-Munoz et al., 2017). Comparative studies of modern taxa have successfully elucidated patterns of variation across mammal clades reflecting dietary specialisations (Evans et al., 2007; Lazzari et al., 2008; Pineda-Munoz et al., 2017; Selig et al., 2019). General predictive patterns also posit ecological and evolutionary drivers for common mammalian patterns (Selig et al., 2021). Dental topography has also been an informative tool to reconstruct the diet of extinct mammals and to discover and document macroecological patterns (López-Torres et al., 2018; Prufrock et al., 2016; Selig et al., 2020; Ungar, 2004). In bats, DTA has only been applied in two studies (López-Aguirre et al., 2021b; Santana et al., 2011). Santana et al. (2011) explored differences between bat insectivores, omnivores and herbivores, finding higher topographic complexity in frugivores. Comparisons of values of DTA in modern bats to reconstruct the diet of the fossil species *Notonycteris magdalenensis* suggests DTA can reflect the diet of bats (López-Aguirre et al., 2021b). These studies revealed an increase in dental complexity and sharpness in frugivorous noctilionoids and a decrease in species with liquid diets (i.e. nectarivory and sanguivory). However, broader patterns that indicate shared pathways of dental morphological adaptation to exploit different dietary resources largely remain to be explored in bats.

In this study we aimed to trace the connection between dental morphology and diet in noctilionoid bats. Using a battery of DTA metrics, we quantified aspects of the dental morphology for 110 noctilionoid species representing the range and multidimensionality of dietary adaptations. Three-dimensional models of lower first molars were analysed within a multivariate statistical framework using phylogenetic comparative methods, accounting for the effect of evolutionary kinship and body mass in our data. We represented diet in three dimensions (guild, composition and breadth) and tested how dental morphology is related to each parameter. We hypothesised that 1) bat dental morphology does not have strong phylogenetic structuring, reflecting the multiple evolution of various noctilionoid lineages to occupy the same dietary guilds (Datzmann et al., 2010); 2) noctilionoid bats show a significant effect of body mass on dental morphology, following a general mammalian pattern in molar size (Billet & Bardin, 2021); 3) topographic complexity and sharpness will be high in frugivores to optimise oral food handling and compensate reduced forelimb handling; and 4) dietary composition will reveal patterns of variation in the DTA metrics that reflect the level of intensity with which species exploit different food items. In particular, we predicted that we would find higher values of topographic complexity and sharpness in animalivore and frugivore diets and higher crown height and slope in species with liquid diets. Given that most species are known to feed on a variety of resources (despite extreme diet-based morphological specialisations), we did not expect to see a pattern in DTA in response to dietary breadth.

## Methods

### Sample composition

Our sample consisted of 90 specimens representing 85 noctilionoid species (Supplementary Table 1). Data for an additional 25 species were retrieved from the literature to complete a total of 115 samples and 110 species (López-Aguirre et al., 2021b). Species were represented by one to five specimens, with average values being used in analyses for species represented by more than one specimen. We sampled all noctilionoid families (Furipteridae, Mormoopidae, Mystacinidae, Myzopodidae, Noctilionidae, Phyllostomidae and Thyropteridae), 44% of currently recognized noctilionoid species and the entire dietary range found in Chiroptera (Fig. 1). The majority of μCT scans of specimens were retrieved from the Digitizing Extant Bat Diversity Project (Shi et al., 2018) and more generally from MorphoSource. To complement our sampling of rare lineages (e.g. Myzopodidae), additional specimens were sourced from Digimorph (Supplementary Table 1). Data we retrieved from the literature were extracted from specimens that were also sourced from (Shi et al., 2018), ensuring that the imaging protocols were generally consistent across our sample. Due to the relatively thin enamel layer in bats’ molars (Dumont, 1995), only specimens with no dentine exposed in occlusal view were selected, excluding specimens with significant wear. Each specimen was represented by a single lower left first molar (m1). Analysing single teeth provides a better ecomorphological signal (Berthaume et al., 2020), as analysing multiple teeth insert other sources of variation (e.g. variation in dental formulae and differential wear patterns across teeth) that obscure diet-based patterns. Also, lower first molars have been proposed as an effective proxy to test dietary diversity in bats based on DTA. Freeware 3D Slicer was used to process raw scan data and to segment individual m1s (Fedorov et al., 2012). Meshlab was used to process 3D meshes of individual m1s (Cignoni et al., 2008), cropping each crown below the cingulid (following established protocols for DTA in mammals and bats) (Berthaume et al., 2019b). To standardise our 3D models, individual meshes were simplified to 10,000 faces and smoothed using 3 iterations of the Laplacian Smoothing filter in Meshlab (Berthaume et al., 2019b).

**Figure 1.**
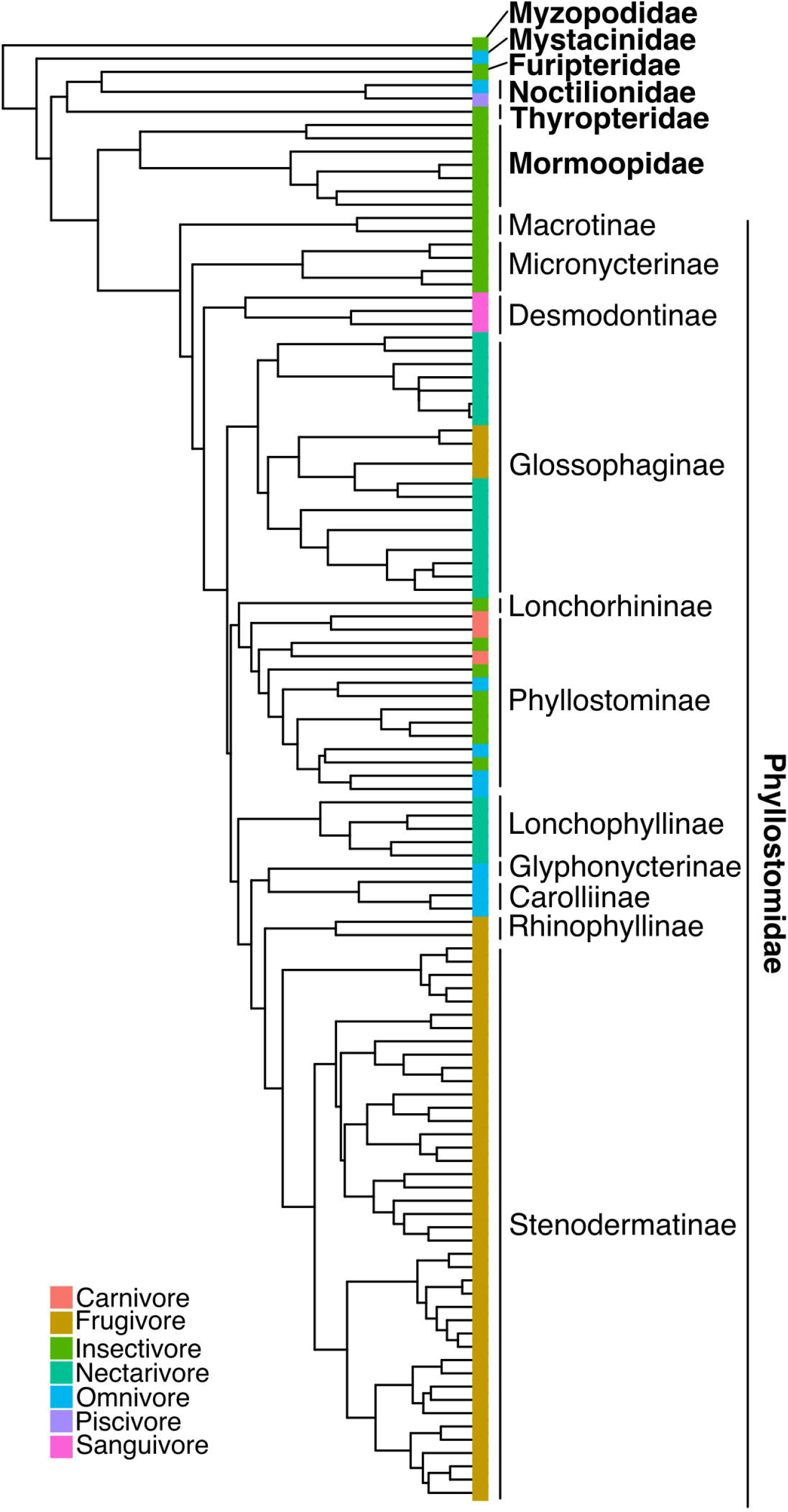
Evolutionary relationships of species sampled, based on a modified version of Shi & Rabosky (2015)’s phylogeny. Tip colours indicate dietary guilds.

### Dental morphology quantification

Data for five DTA metrics were collected (Fig. 2): Dirichlet Normal Energy (DNE, Bunn et al., 2011), Orientated Patch Count Rotated (OPCR, Evans et al., 2007), Relief Index (RFI, Boyer, 2008), slope (Ungar & M’Kirera, 2003) and molar area (MA). Dirichlet Normal Energy uses changes in normal vectors generated for individual faces in the mesh to quantify curvature of the crown as proxy for sharpness (Bunn et al., 2011). Higher values of DNE are usually associated with greater shearing capacity and animalivory in mammals (Ledogar et al., 2013, 2018; López-Torres et al., 2018; Selig et al., 2021). Oriented Patch Count Rotated estimates surface complexity by quantifying the number of clusters of faces (i.e. patches) in the mesh that are similarly oriented in space (Evans et al., 2007). High OPCR values are interpreted as a representing a greater set of surfaces for processing mechanically resistant foodstuff (e.g. folivory and animalivory, Evans et al., 2007). Relief Index is the ratio of 3D-to-2D area of the tooth crown, representing an index of crown height (Allen et al., 2015; Boyer, 2008; Cooke, 2011). Low RFI values associated with flat tooth crowns have been found in mammalian frugivores and high RFI values with tall cusps in folivores and animalivores (Pampush, Spradley, et al., 2016). Slope is the mean slope across the surface in occlusal view, averaging the slope of all faces of the mesh (Ungar & M’Kirera, 2003). Higher slopes are usually interpreted as more elevated crowns, indicating higher or pointier crowns (Berthaume et al., 2020). Molar area was estimated as the 2D surface area of a mesh. All DTA metrics were calculated using the R package *molaR* (Pampush, Winchester, et al., 2016). To quantify OPCR, we used a minimum number of faces of three, iterating the process eight times, with a rotation of 5.625 degrees between iterations.

**Figure 2.**
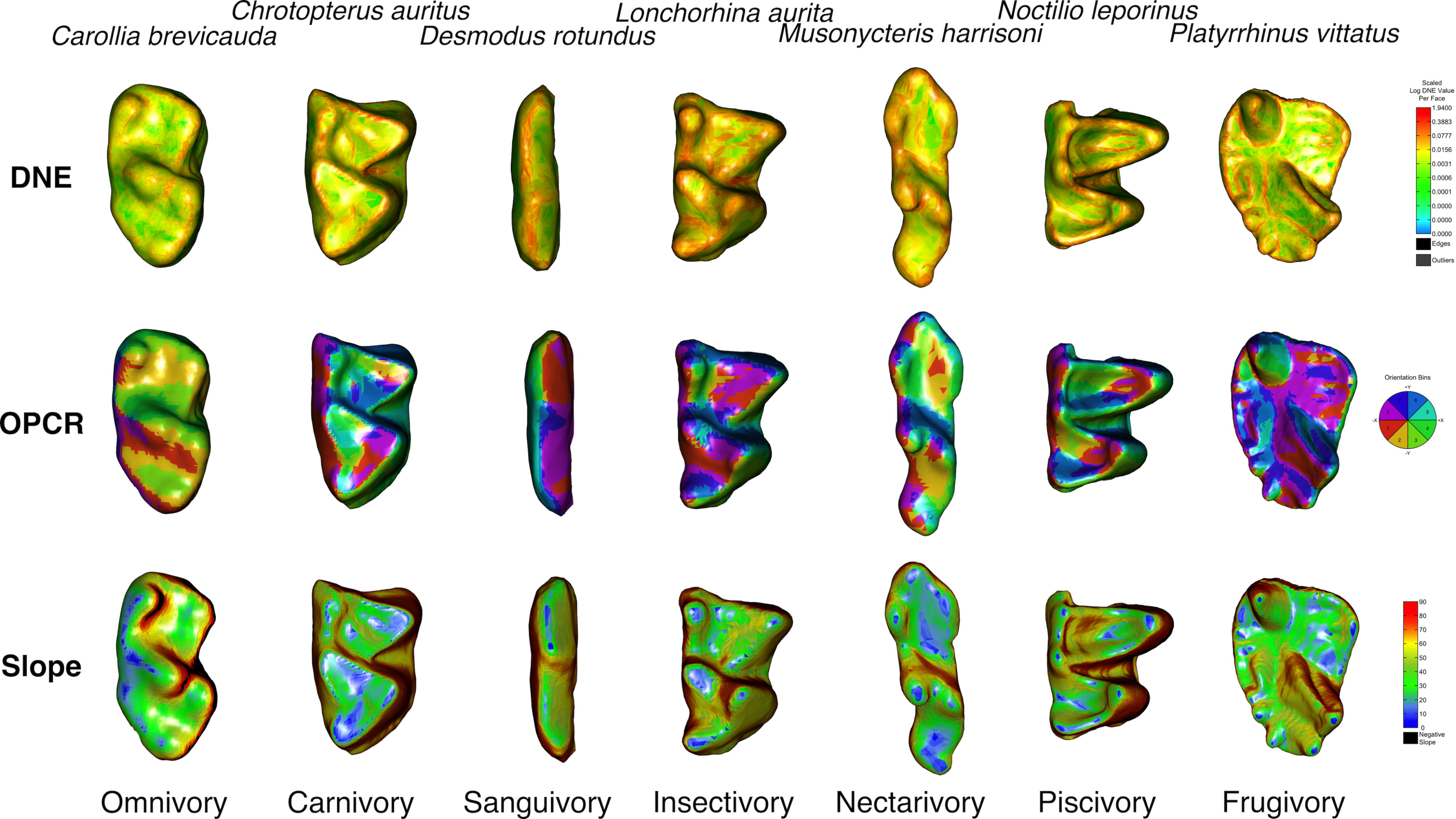
Diversity of dental morphology across the dietary range found in Noctilionoidea, exemplified with DTA metrics; DNE, OPCR and slope. 3D meshes not to scale. Low DNE and slope values in blue and high values in red. For 3D-OPCR, differences in colour relate to the orientation of patches with respect to the 8 cardinal directions.

### Body mass and dietary data acquisition

Body mass of species was compiled from the literature and averages were used in further studies (Giannini et al., 2020; Moyers Arévalo et al., 2018). To capture the dietary diversity of noctilionoid bats, we quantified three dimensions of dietary specialisation: 1) guild, 2) composition and 3) breadth. Species were assigned to one of seven dietary guilds previously used to study dietary adaptations in bats (e.g. by Arbour et al., 2019; Monteiro & Nogueira, 2011; Rossoni et al., 2019), grouping species based on broad dietary similarities: carnivory, frugivory, insectivory, nectarivory, omnivory, piscivory and sanguivory. Comparing dietary guilds allowed us to elucidate diet-based general patterns, contrasting all our species within a single framework. We also gathered data on dietary composition and dietary breadth. For dietary composition, food items were classified into seven categories: arthropods, blood, terrestrial vertebrates, fish, flower pieces, pollen and nectar, fruit and seeds (Rojas et al., 2011, 2012, 2018). For each species, the relative importance of each food item was codified as either: absent (one), complementary (two), predominant (three), or exclusive (four), following Rojas et al. (2011, 2012, 2018). Quantifying diet composition of each species and the relative importance of each item allows the comparison of species across a range of intensity of exploitation of a given resource. Finally, dietary breath was compiled as the marginality index developed by (Rojas et al., 2018) based on the diet composition data described above. This marginality index places species within a range of values between zero and one, representing a gradient of specialisation from herbivory to carnivory. Species with values close to zero indicate a specialised herbivore diet, values close to one indicate a specialised animalivore diet, whereas species with values close to 0.5 represent generalist species capable of both herbivory and animalivory. Marginality calculates diet as a continuous numerical variable that can be included as a factor in statistical analyses that can facilitate the reconstruction of an ecological transition from strict herbivory to strict animalivory. A caveat with this approach is that it obscures dietary subguilds at the extremes of the range of marginality values (e.g. frugivory and nectarivory within herbivory). Data for dietary composition and breadth were retrieved from (Rojas et al., 2011, 2012, 2018), whereas dietary guild data were retrieved from (Monteiro & Nogueira, 2011; Rossoni et al., 2019).

### Statistical analyses

We used the phylogeny of (Shi & Rabosky, 2015) to test the effect of evolutionary kinship in our data, pruning the phylogeny to match our sample composition. We tested for the presence of phylogenetic signal using the K−statistic implemented in the “phylosig” function in the R package *phytools* (Revell, 2012). Before testing the effect of body mass on dental topography, we first tested for multicollinearity between body mass and diet using a phylogenetic linear model (log(BM)∼Marginality). A weak but significant correlation was found, marginality explaining less than 4% of variation in body mass (*R*^2^ = 0.031, *P*= 0.035). In light of this result, all further analyses were performed with log-transformed body mass. Phylogenetic ANOVAs were used to test for statistical differences in DTA metrics across dietary guilds using the “procD.pgls” function in the R package *Geomorph* (Adams et al., 2013), including body mass as covariate (log(DTA)∼Diet+ log(BM)). Post hoc pairwise tests were used to further explore differences between dietary guilds, using the “pairwise” function in the R package *RRPP*.

Piscivory was excluded due to small sample size (n=1). Boxplots were generated to visually explore patterns of variation in DTA metrics across guilds. A principal components analysis was performed on the correlation matrix of DTA metrics, to explore multivariate patterns of variation while reducing the dimensionality of our data. Additionally, we performed a phylogenetic principal component analysis to account for the phylogenetic structuring (i.e. significant phylogenetic signal) of our data, using the “phyl.pca” function in *phytools* (Revell, 2012). We calculated the dental topographic disparity (quantified as the sum of variances of the standardised DTA metrics) per guild using the R package *dispRity* (Guillerme, 2018). We tested whether interguild differences in sample size influenced disparity metrics, finding no association between them (R^2^ = -0.04843, *P*= 0.43). A Wilcoxon test was applied for pairwise comparisons across guilds with a Bonferroni correction.

Next, the association between diet composition and dental morphology was assessed. Dietary composition (measured as the relative importance of a given resource) allowed us to test how DTA metrics may differentially correlate with specialisation for feeding on a particular type of resource. We only studied diets that had different levels of exploitation across species (i.e. carnivory, insectivory, frugivory and nectarivory). Phylogenetic regression models were used to quantify the relative association of increasing exploitation of a given resource to variation in each DTA metric. We included body mass as a covariate to estimate the relative effect of size in each model. Barplots were used to visualise the relative importance of each DTA metric for the increasing exploitation of a resource. Finally, we analysed the influence of dietary breadth on dental morphology. Phylogenetic regression models were used to discern the relative effect of body mass and dietary breadth (marginality) on each DTA metric (log(DTA)∼log(BM)+Marginality), using the “procD.pgls” function in *Geomorph* (Adams et al., 2013). Biplots were used to visualise patterns of association between DTA metrics and marginality.

## Results

### Differences across dietary guilds

A significant phylogenetic signal was detected for all five DTA metrics, OPCR showing the weakest signal and RFI the strongest (Table 1). Boxplots revealed differences in DTA metrics between dietary guilds (Fig. 3 left column). Dirichlet Normal Energy was high in piscivorous, frugivorous and insectivorous species and lower in sanguivorous species, all guilds overlapping with at least another guild (Fig. 3). A similar pattern was found in OPCR values, with the piscivore showing similar values to carnivores, frugivores and insectivores, and with lower values found in species with liquid diets (Fig. 3). In contrast, sanguivorous and piscivorous species showed the highest values of RFI and slope, whereas the lowest values for those two metrics were found in frugivorous species, followed by nectarivorous species (Fig. 3). Molar area was noticeably greater in carnivorous and piscivorous species and lower in species with liquid diets (Fig. 3).

**Figure 3.**
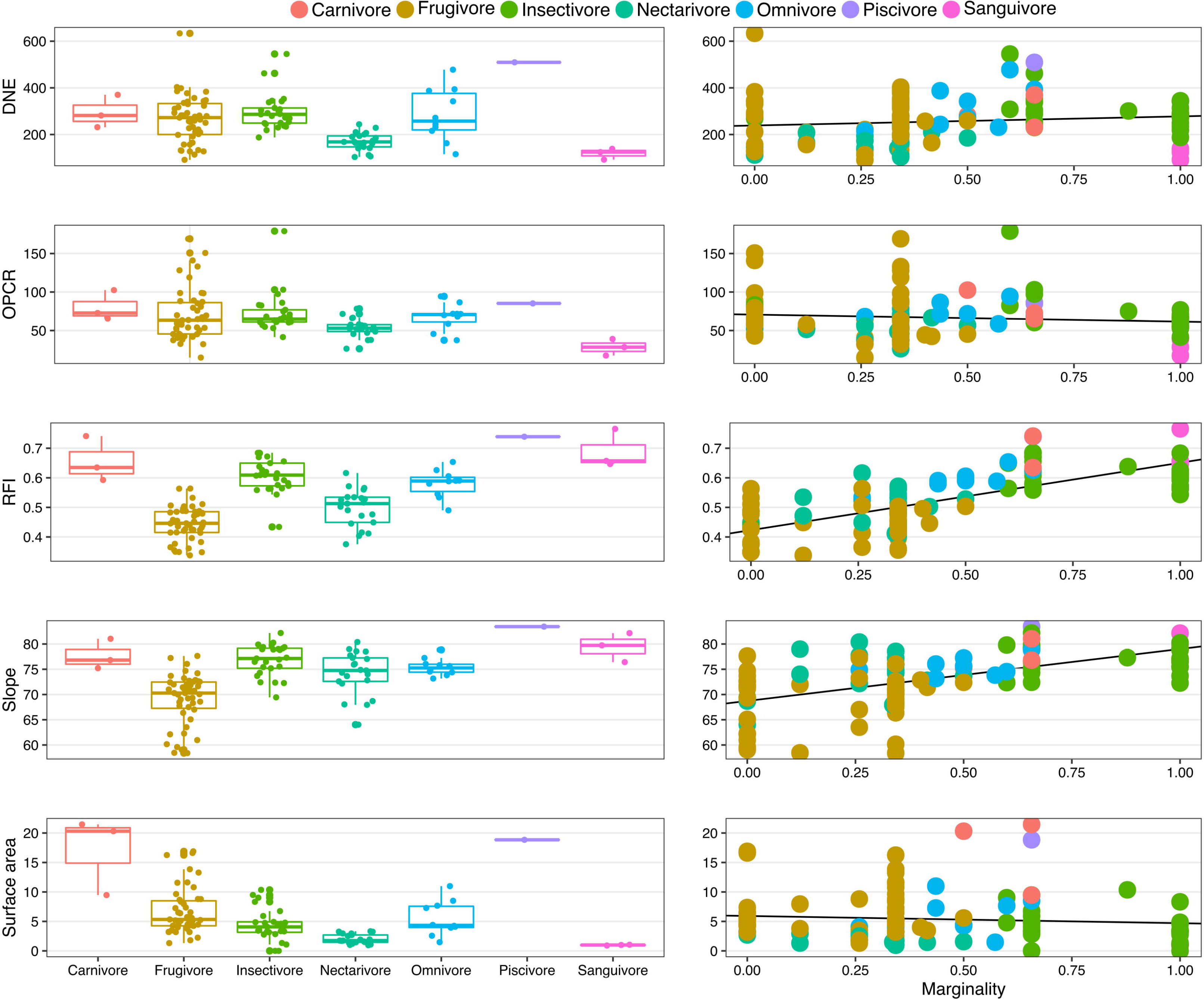
Variation in dental complexity across Noctilionoidea based on diet. Boxplots comparing DTA metrics across different dietary guilds (left column), and scatterplots showing linear regressions of DTA metrics against marginality (i.e. dietary breadth). Marginality values close to zero represent specialist herbivorous diets, whereas values close to one represent specialist animalivorous diets.

**Table 1.** Statistical test of phylogenetic signal on each DTA metric, based on 10000 iterations.

|  | *K*- | *P* |
| --- | --- | --- |
| DNE | 0.513 | <0.0001 |
| OPCR | 0.299 | 0.049 |
| RFI | 1.553 | 0.0001 |
| Slope | 0.848 | 0.0001 |
| Molar size | 0.754 | 0.0002 |

Phylogenetic ANOVAs indicated statistically significant differences between dietary guilds across all DTA metrics, whereas body mass did not have a significant effect on RFI (Table 2). Comparing the amount of variation explained by each variable, body mass explained on average 17% of variation, whereas differences in dietary guilds explained 28.26% of DTA variation. Pairwise comparisons revealed slope and RFI in frugivorous and nectarivorous species were significantly different from all other guilds, whereas OPCR, DNE and MA in species with liquid diets were significantly different from other guilds with solid diets (Supplementary Table 2).

**Table 2.** Phylogenetic ANOVAs of differences in DTA metrics between dietary guilds, based on 10,000 iterations.

|  |  | Df | SS | MS | Rsq | F | Z | *P* |
| --- | --- | --- | --- | --- | --- | --- | --- | --- |
| DNE | log(BM) | 1 | 0.1026 | 0.1025 | 0.0619 | 8.7448 | 2.3132 | 0.0062 |
|  | Diet | 5 | 0.3585 | 0.0717 | 0.2163 | 6.1139 | 3.193 | 0.0011 |
| OPCR | log(BM) | 1 | 0.0752 | 0.0752 | 0.0602 | 7.7843 | 2.2294 | 0.0089 |
|  | Diet | 5 | 0.1890 | 0.0378 | 0.1512 | 3.9125 | 2.582 | 0.0051 |
| RFI | log(BM) | 1 | 0.0010 | 0.0010 | 0.0055 | 1.0734 | 0.561 | 0.3066 |
|  | Diet | 5 | 0.0875 | 0.0175 | 0.4699 | 18.2758 | 7.3422 | <0.0001 |
| Slope | log(BM) | 1 | 0.0020 | 0.0020 | 0.0643 | 10.1131 | 2.5044 | 0.0021 |
|  | Diet | 5 | 0.0091 | 0.0018 | 0.2877 | 9.0581 | 4.9303 | <0.0001 |
| MA | log(BM) | 1 | 2.1707 | 2.1707 | 0.4940 | 230.98 | 7.2213 | <0.0001 |
|  | Diet | 5 | 1.2651 | 0.2530 | 0.2879 | 26.923 | 8.2636 | <0.0001 |

Dental topographic disparity was found to be higher in noctilionoid frugivores and lower in nectarivores and sanguivores (Fig. 4). Wilcoxon pairwise comparisons showed statistical differences in disparity between all guilds but not between carnivores, insectivores and omnivores (Table 3).

**Figure 4.**
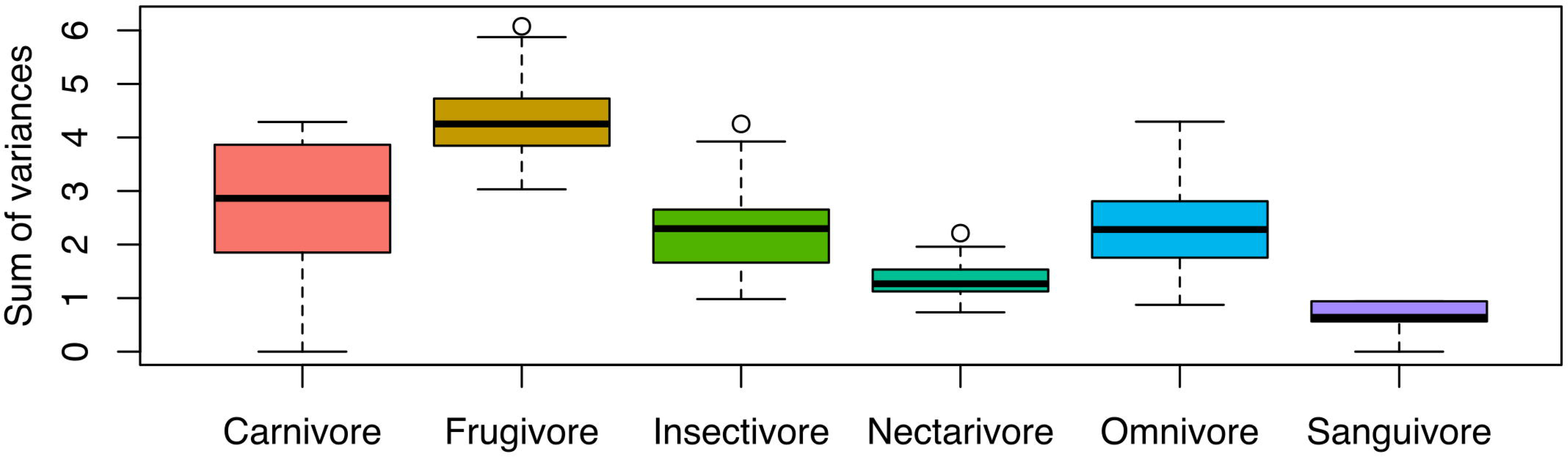
Boxplots of dental topographic disparity (quantified as the sum of variances across DTA metrics) across dietary guilds, based on 100 resampling iterations.

**Table 3.** Wilcoxon test of pairwise comparisons of dental disparity across noctilionoid dietary guilds.

|  | W statistic | P value |
| --- | --- | --- |
| Carnivore : Frugivore | 1494 | <0.0001 |
| Carnivore : Insectivore | 5672 | 1.0000 |
| Carnivore : Nectarivore | 8455 | <0.0001 |
| Carnivore : Omnivore | 5864 | 0.5143 |
| Carnivore : Sanguivore | 8698 | <0.0001 |
| Frugivore : Insectivore | 9338 | <0.0001 |
| Frugivore : Nectarivore | 10000 | <0.0001 |
| Frugivore : Omnivore | 9774 | <0.0001 |
| Frugivore : Sanguivore | 10000 | <0.0001 |
| Insectivore : Nectarivore | 8818 | <0.0001 |
| Insectivore : Omnivore | 5415 | 1.0000 |
| Insectivore : Sanguivore | 9979 | <0.0001 |
| Nectarivore : Omnivore | 643 | <0.0001 |
| Nectarivore : Sanguivore | 9732 | <0.0001 |
| Omnivore : Sanguivore | 10000 | <0.0001 |

The first two principal components of dental morphospace (based on principal component analysis) explained 83.79% of variation (Fig. 5 left). Piscivorous and sanguivorous species were the only taxa occupying non-overlapping subspaces, whereas frugivores showed the greatest dispersion across morphospace and the greatest overlap with other guilds. The guilds with the greatest diversity (i.e. insectivory, omnivory and frugivory) tended to overlap towards the origin of morphospace. Low values of Principal Component 1 were correlated with high values of OPCR and DNE, separating species that feed on vertebrates (i.e. piscivory and carnivory) from those with liquid diets. High values of Principal Component 2 were associated with low values of slope and RFI. A similar pattern was found in phylogenetic morphospace (based on phylogenetic principal component analysis), with the first two components explaining 78.52% of variation (Fig. 5 right) and the same pattern of correlation with the various DTA metrics.

**Figure 5.**
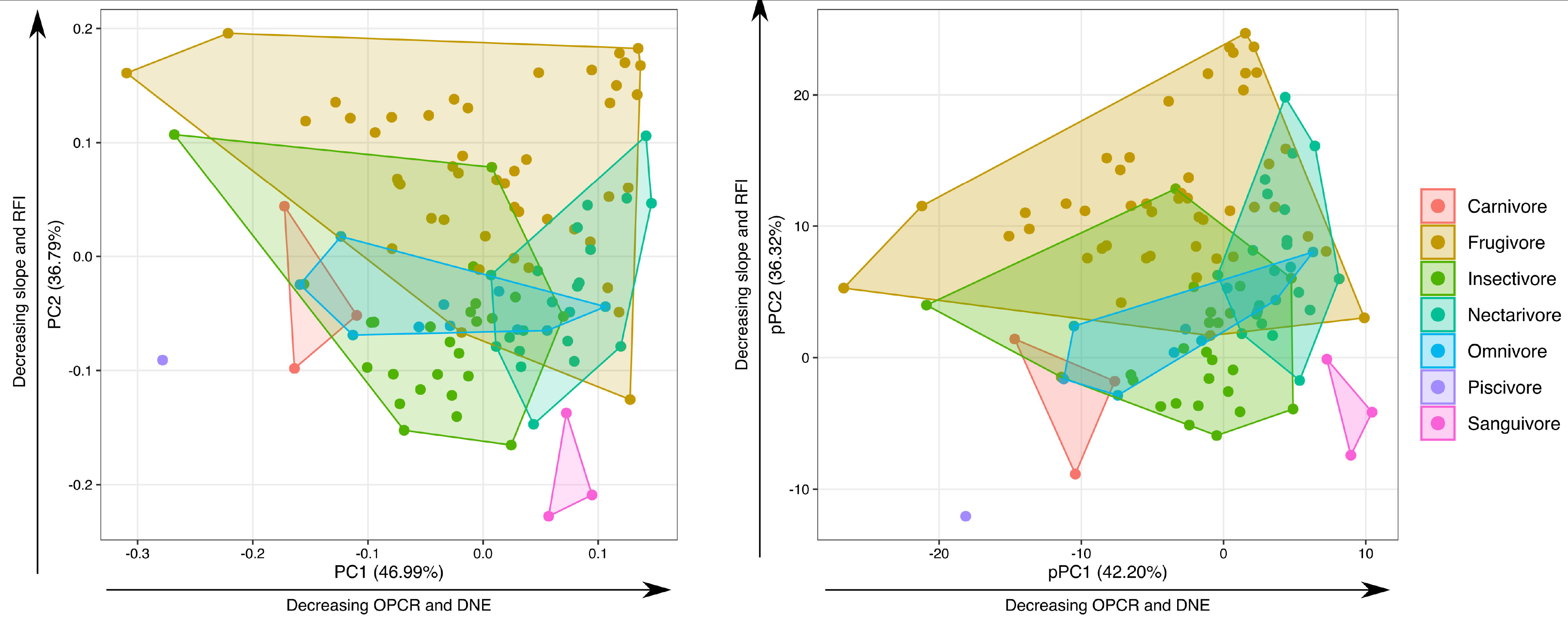
Dental morphospace (based on principal component analysis) and phylogenetic morphospace (based on phylogenetic principal component analysis).

### Dietary composition and level of specialisation

Comparing dental morphology along trajectories of increasing importance of a given resource in a species’ diet showed decoupled patterns across DTA metrics (Fig. 6). Mean DNE values increased with the increasing importance of carnivory, frugivory and insectivory, and mean values decreased as nectarivory increased. Values of OPCR increased as degrees of carnivory and frugivory increased, remained stable across varying relative amounts of insectivory and values decreased as nectarivory increased. Values of RFI and slope increased as amounts of carnivory and insectivory increased and both values and slope decreased as degree of frugivory and nectarivory decreased. Molar area showed a dramatic increase with increasing levels of carnivory, remained relatively stable for differing degrees of frugivory and insectivory, and decreased with increasing nectarivory. Intensifying frugivory explained the highest percentage of variation in OPCR, RFI and slope, while body mass explained the highest percentage of variation in MA and DNE (Fig. 7). Insectivory and nectarivory explained the lowest amount of variation in DTA metrics (2.64% and 1.44%, respectively). Frugivory and body mass had the highest explicatory power across all our models, explaining on average 17.11% and 16.33% of DTA variation. Intensifying carnivory was the dietary adaptation that explained the highest amount of variation in MA (9.91%). Phylogenetic regression models confirmed the diverging association of different DTA metrics with specialisation for different diets, frugivory and body mass being the only statistically significant factors in all models (Supplementary Table 3).

**Figure 6.**
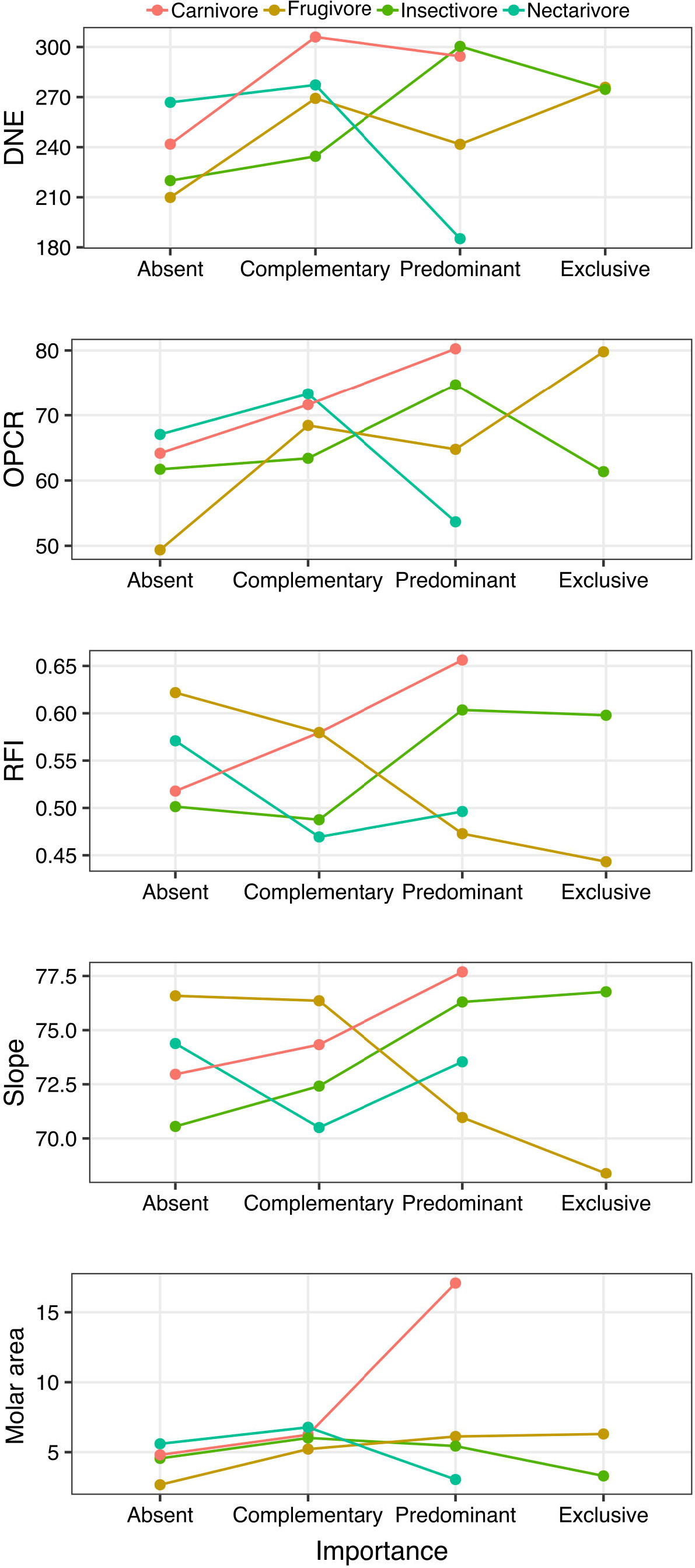
Trajectories of mean values of DTA metrics along a gradient of increasing intensity of exploitation of a given dietary resource in noctilionoid bats.

### Effect of dietary breadth

Phylogenetic linear models revealed a significant effect of body mass and marginality (degree of herbivory vs animalivory) on all DTA metrics, with body mass explaining on average 19.94% of variation and marginality explaining 20.37% (Table 4). Body mass correlated more strongly with MA (explaining 38.86% of variation), whereas marginality correlated more strongly with RFI and slope (explaining 39.91% and 25.74% of variation, respectively). Slope and BM did not significantly correlate with body mass and marginality, respectively. Biplots show the strong correspondence between RFI and slope and marginality, with specialist herbivores having the lowest values of both metrics and specialist animalivores the highest values (Fig. 3 right column). RFI and slope showed the strongest interaction between marginality and dental morphology, both showing a positive interaction.

**Table 4.** Phylogenetic linear regression models of the effect of body mass (BM) and dietary breadth (marginality) on DTA metrics, based on 10,000 iterations.

|  |  | df | SS | MS | R<sup>2</sup> | F | Z | P |
| --- | --- | --- | --- | --- | --- | --- | --- | --- |
| DNE | log(BM) | 1 | 0.30209 | 0.302091 | 0.18349 | 25.1161 | 1.9893 | 0.0001 |
|  | Marginality | 1 | 0.05727 | 0.057273 | 0.03479 | 4.7617 | 1.27 | 0.0296 |
| OPCR | log(BM) | 1 | 0.25076 | 0.25076 | 0.16837 | 25.452 | 2.0276 | 0.0001 |
|  | Marginality | 1 | 0.18435 | 0.184348 | 0.12378 | 18.711 | 1.8726 | 0.0001 |
| RFI | log(BM) | 1 | 0.017814 | 0.017814 | 0.05721 | 11.261 | 1.6139 | 0.0017 |
|  | Marginality | 1 | 0.124275 | 0.124275 | 0.39914 | 78.558 | 2.5358 | 0.0001 |
| Slope | log(BM) | 1 | 0.000624 | 0.0006242 | 0.01215 | 1.7798 | 0.8305 | 0.1829 |
|  | Marginality | 1 | 0.013224 | 0.0132238 | 0.25741 | 37.7071 | 2.211 | 0.0001 |
| MA | log(BM) | 1 | 3.1885 | 3.1885 | 0.3886 | 69.4906 | 2.4991 | 0.0001 |
|  | Marginality | 1 | 0.107 | 0.107 | 0.01304 | 2.3321 | 0.9646 | 0.1255 |

**Figure 7.**
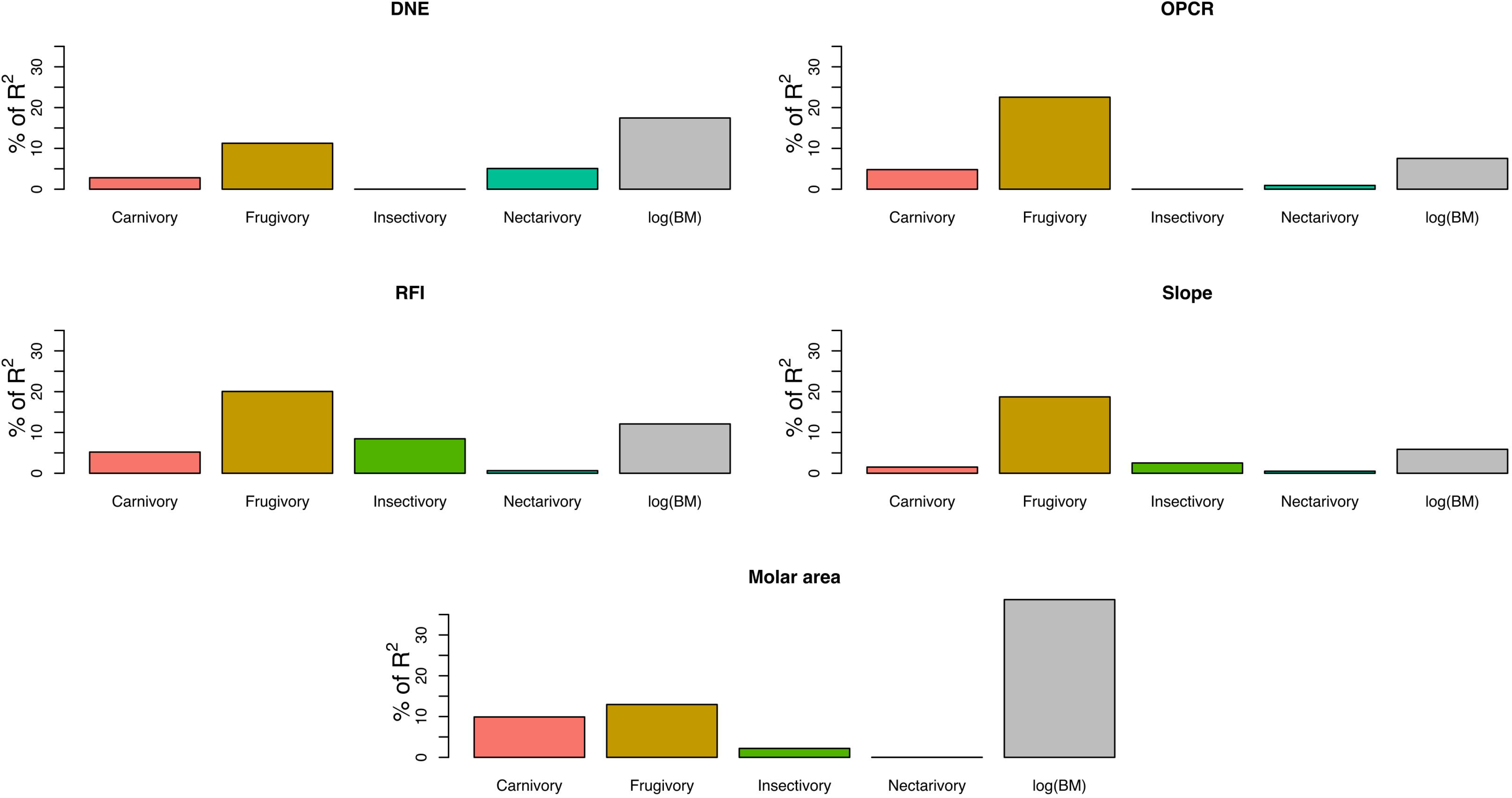
Relative importance of increasing intensity of exploitation of various dietary resources on linear regression models of dental morphological variation, expressed as the percentage of R^2^ of the model explained by each DTA metric.

## Discussion

Multivariate DTA strongly supports a functional link between dental phenotypic variation and multiple aspects of diet in Noctilionoidea. Statistically significant effect of size and phylogenetic signal indicate dental morphology is also influenced by body mass and evolutionary relatedness, contrary to our prediction of a lack of phylogenetic structuring. After controlling for phylogenetic structuring and size, retrieved patterns of DTA variation showed differential effects of dietary guild, composition and breadth, suggesting there are different pathways for dental morphology to correlate with diet. Differences in DTA metrics across dietary guilds reflect physical differences between diets (e.g. liquid diets vs. solid diets). High OPCR and DNE in frugivorous species supported our hypothesis of increased complexity and sharpness to optimise handling mechanically demanding foodstuff. Changes in the intensity of frugivory explained the highest amount of variance across DTA metrics, showing the overarching effect that frugivory had for the morphological diversification in Noctilionoidea. Crown height and occlusal slope showed the most significant associations with dietary breadth, both decreasing as diets narrow to specialise for herbivory and increasing towards narrow animalivorous diets.

### Differences between dietary guilds

Consistent with previous studies of DTA in bats, we found a significant phylogenetic and ecological signal in dental morphology, reflecting both dietary specialisations and evolutionary relatedness (López-Aguirre et al., 2021b). Body mass had a differential effect across DTA metrics, always explaining less variation than diet (excepting in MA). Differences in dental morphology across noctilionoid bats revealed biomechanical similarities in foodstuff processing within dietary guilds, following patterns previously reported in other bats and mammals (López-Aguirre et al., 2021b; Pineda-Munoz et al., 2017; Santana et al., 2011; Winchester et al., 2014). General patterns in diet-based variation in DNE among mammals typically suggest a negative correlation between DNE and frugivory, and an increase in DNE in omnivorous and insectivorous taxa (Berthaume et al., 2019a; Fulwood et al.,2021; Winchester et al., 2014). Our finding that values of DNE are instead broadly similar in frugivorous, insectivorous and omnivorous noctilionoid bats confirms previous findings from a smaller sample of species (López-Aguirre et al., 2021b), indicating a clear deviation from the general mammalian pattern in these bats (Fulwood et al., 2021; Pérez-Ramos et al., 2020; Winchester et al., 2014). This suggests a unique pattern of morphological adaptation to frugivory in noctilionoid bats, at least in the dentition. Future studies could better elucidate adaptations to frugivory in bats by comparing phyllostomid and pteropodid bats.

Dirichlet Normal Energy has usually been interpreted as a measure of tooth sharpness, so our findings suggest frugivorous noctilionoids have sharper molars than the average frugivorous mammal. We hypothesise that the reduced capacity that bats have to handle fruits with their forelimbs and the fact they feed either perched upside down or during flight represented a selective pressure for sharper molars to optimise food processing, due to a reduced potential for pre-ingestion foodstuff processing (Aguirre et al., 2003; Santana et al., 2011; Santana & Dumont, 2009). However, to our knowledge our study is only the second one to investigate DNE in bats, so further studies are required to determine if this is a chiropteran feature or something specific to noctilionoids.

Higher dental complexity has been proposed as a common adaptation for frugivory in bats (López-Aguirre et al., 2021b; Santana et al., 2011), contrary to recent studies that have found an opposite pattern in ursid carnivorans and strepsirrhine primates (Fulwood et al., 2021; Pérez-Ramos et al., 2020). Our results support the general chiropteran pattern of higher dental complexity in frugivores, as indicated by our results of OPCR in noctilionoid bats. Values of RFI increased on the scale from herbivory to animalivory, similar to previous findings in primates. Values calculated for RFI were higher in nectarivorous bats than in frugivorous bats, which would seem to be in opposition to the hypothesis that high RFI necessarily means more mastication surface area by Pineda-Munoz et al. (2017). Additionally, we found high slope and RFI in species with liquid diets, confirming previous findings in bats (López-Aguirre et al., 2021b). Reduction of size in postcanine dentition has been proposed as a common adaptation in mammals specialised to different liquid diets (e.g. sanguivory and exudativory; Burrows et al., 2020). Compared to gummivore and exudativore primates (Burrows et al., 2020; Burrows & Nash, 2010; Pineda-Munoz et al., 2017), noctilionoid bats with liquid diets have laterally compressed molars, which reduces the 2D area of the crown and inflates RFI and slope (see Fig. 2). Inflated RFI and slope revealed in our results could indicate a limitation in the biomechanical interpretations of these metrics and should be accounted for in future studies.

In terms of dental disparity, the significantly greater dental variability in frugivorous noctilionoid bats showcases the extreme phenotypic diversity found associated with frugivory, reflecting previous findings supporting the importance that shifts to feeding on fruit apparently had on morphological and taxonomic diversification of bats (Arbour et al., 2019; Monteiro & Nogueira, 2011; Rojas et al., 2012; Rossoni et al., 2019). Overall, our results reveal noctilionoid bats do not follow the general mammalian pattern of OPCR variation across dietary guilds. Further, DNE in noctilionoid frugivores and RFI and slope in species with liquid diets showed deviations from the previous findings in mammals.

### DTA and dietary composition

Patterns of DTA based on the intensity of exploitation of different resources supported our hypothesis that dietary composition, in addition to putative guild, has an effect on dental morphology. Increasing values of DNE in species that rely more on fruit supports previous findings that specialist frugivorous noctilionoids tend to have sharper molars than facultative frugivores, possibly to optimise food handling (López-Aguirre et al., 2021b). This seems to be correlated with the evolution of a labial rim on the molars, formed by the displacement of the paracone and metacone from a central position in the occlusal surface of the molar to the labial edge of the tooth, a trait which is found in specialised frugivore noctilionoids (Freeman, 1988). Decreasing values of DNE associated with increasing amounts of nectarivory also support our prediction that species with liquid diets have lower biomechanical needs to break down or manipulate foodstuff (Freeman, 1995). Only slope did not have a tendency to increase with intensified carnivory, suggesting that evolution of specialized carnivory represents a multifaceted adaptation process, rather than encompassing change in a single dimension of dental morphology that facilitated the colonisation of this dietary niche (Freeman, 1988; Norberg & Fenton, 1988; Santana & Cheung, 2016). Differences in the association between DTA metrics and intensifying exploitation of different resources showed various patterns of ecomorphological adaptation. Intensifying carnivory explaining the greatest proportion of variance of molar area reflects previous findings on the importance of increased body size for the adoption of carnivory in Chiroptera (Freeman, 1988; Norberg & Fenton, 1988; Santana & Cheung, 2016). It has been suggested that increased body size was the primary adaption required for carnivory during the evolution of noctilionoid bats (Freeman, 1988; Giannini et al., 2020; Norberg & Fenton, 1988), predating other dental or skeletal adaptations (López-Aguirre et al., 2021b). Frugivory had the greatest explicatory power across all DTA metrics, revealing the overarching importance of frugivory for the evolution of noctilionoid dentition (Freeman, 1981, 1988, 1995). Coupled with our findings of higher dental morphological disparity in frugivores, our models indicate frugivory was a major factor during noctilionoid dental diversification, mirroring previous studies and reflecting the overarching importance of frugivory for the morphological diversification of Noctilionoidea (Arbour et al., 2019; Monteiro & Nogueira, 2011; Rossoni et al., 2017, 2019; Santana et al., 2012). The low explicatory power of insectivory across DTA metrics might indicate the ancestral insectivorous diet of the superfamily and its lower importance to promote morphological variability (Rossoni et al., 2019). It is interesting that intensifying nectarivory only explained variation in DNE, suggesting that reduction in molar sharpness played a major role during the adoption of nectarivory in Noctilionoidea.

### DTA and dietary breadth

Contrary to our prediction, all DTA metrics with the exception of molar area showed a significant association with dietary breadth, revealing that the diversity in species’ diets is also reflected in dental morphology. Nevertheless, the variation in the strength and direction of the association across DTA metrics indicates different pathways shaping phenotypic adaptions to generalist and specialist diets. The stronger linear correlation between slope, RFI and marginality points to a common path of specialisation for herbivory on the one hand and animalivory on the other. The pattern of values of slope and RFI decreasing in species with narrow herbivorous diets and increasing in those that have narrow animalivorous diets underscore previous findings regarding dietary guilds and diet composition (López-Aguirre et al., 2021b). Wider molar occlusal surfaces are commonly found in frugivorous noctilionoids (particularly in the subfamily Stenodermatinae, comprised largely of specialised frugivores), whereas taller and steeper molar crowns are characteristic of animalivorous bats (Freeman, 1981, 1988, 1995; Santana & Cheung, 2016). Biplots of marginality and DNE, OPCR and MA suggest higher variation of these three metrics in species with broader diets (i.e. generalist omnivores), explaining the weaker linear correlation between the three metrics and marginality. More omnivorous species face the challenge of consuming a wider range of foodstuffs that present a myriad of biomechanical challenges (Aguirre et al., 2003; Santana et al., 2011; Santana & Dumont, 2009). Higher variation in DNE and OPCR values suggest that omnivory can be linked to greater morphological disparity to accommodate a variety of feeding demands (Rojas et al., 2018). We hypothesise that increased DNE and OPCR equip generalist species with the dental tools necessary to process a variety of foods with different biomechanical properties. Combined, our results suggest that variability in RFI and slope provide evolutionary pathways to specialist diets, whereas increases in DNE and OPCR represent adaptations to omnivory.

## Conclusions

Using phylogenetic comparative methods and multivariate DTA, we deconstructed variation in noctilionoid dental morphology into its phylogenetic, ecological and biological components. Our finding of significant phylogenetic and size-dependent structuring in DTA indicates that dental topography does not respond only to diet-related functional demands, but also to phylogenetic relatedness, highlighting the need to account for multiple sources of variation. Our results suggest frugivorous noctilionoids have sharper molars (higher DNE values) than expected compared with other frugivorous mammals, that species with liquid diets show increased molar crown height and steepness, and that animalivorous noctilionoids have larger molars than herbivorous taxa. We also recovered DTA specialisations to dietary composition, revealing that the intensity of exploitation of a resource also correlates with dental morphology. Molar area was the trait most strongly associated with carnivory, whereas RFI and slope explained the greatest variation in our models of increasing insectivory, frugivory and nectarivory. Finally, generalist noctilionoids were found to have higher DNE and OPCR, whereas the range of RFI and slope values revealed a pathway for the transitions to specialist herbivory and specialist animalivory. Incorporating data on biomechanical properties of food items (e.g. item hardness and size) in futures studies could further elucidate still unexplored patterns of dental variation in bats.

## References

Adams, D. C., Otárola-Castillo, E., & Paradis, E. (2013). Geomorph: An R package for the collection and analysis of geometric morphometric shape data. Methods in Ecology and Evolution, 4(4), 393–399. https://doi.org/10.1111/2041-210x.12035

Aguirre, L. F., Herrel, A., Van Damme, R., & Matthysen, E. (2003). The Implications of Food Hardness for Diet in Bats. Functional Ecology, 17(2), 201–212.

Allen, K. L., Cooke, S. B., Gonzales, L. A., & Kay, R. F. (2015). Dietary Inference from Upper and Lower Molar Morphology in Platyrrhine Primates. PLoS ONE, 10(3), e0118732. https://doi.org/10.1371/journal.pone.0118732

Arbour, J. H., Curtis, A. A., & Santana, S. E. (2019). Signatures of echolocation and dietary ecology in the adaptive evolution of skull shape in bats. Nat Commun, 10(1), 2036. https://doi.org/10.1038/s41467-019-09951-y

Arbour, J. H., Curtis, A. A., & Santana, S. E. (2021). Sensory adaptations reshaped intrinsic factors underlying morphological diversification in bats. BMC Biology, 19(1), 88. https://doi.org/10.1186/s12915-021-01022-3

Arkins, A. M., Winnington, A. P., Anderson, S., & Clout, M. N. (1999). Diet and nectarivorous foraging behaviour of the short-tailed bat (Mystacina tuberculata). Journal of Zoology, 247(2), 183–187. https://doi.org/10.1111/j.1469-7998.1999.tb00982.x

Berthaume, M. A., Lazzari, V., & Guy, F. (2020). The landscape of tooth shape: Over 20 years of dental topography in primates. Evolutionary Anthropology: Issues, News, and Reviews, n/a(n/a). https://doi.org/10.1002/evan.21856

Berthaume, M. A., Winchester, J., & Kupczik, K. (2019a). Ambient occlusion and PCV (portion de ciel visible): A new dental topographic metric and proxy of morphological wear resistance. PLoS ONE, 14(5), e0215436. https://doi.org/10.1371/journal.pone.0215436

Berthaume, M. A., Winchester, J., & Kupczik, K. (2019b). Effects of cropping, smoothing, triangle count, and mesh resolution on 6 dental topographic metrics. PLoS ONE, 14(5), e0216229. https://doi.org/10.1371/journal.pone.0216229

Billet, G., & Bardin, J. (2021). Segmental Series and Size: Clade-Wide Investigation of Molar Proportions Reveals a Major Evolutionary Allometry in the Dentition of Placental Mammals. Systematic Biology, syab007. https://doi.org/10.1093/sysbio/syab007

Boyer, D. (2008). Relief index of second mandibular molars is a correlate of diet among prosimian primates and other euarchontan mammals. Journal of Human Evolution, 55 *6*, 1118–1137.

Brooke, A. P. (1994). Diet of the Fishing Bat, Noctilio leporinus (Chiroptera: Noctilionidae). Journal of Mammalogy, 75(1), 212–218. https://doi.org/10.2307/1382253

Bunn, J. M., Boyer, D. M., Lipman, Y., St Clair, E. M., Jernvall, J., & Daubechies, I. (2011). Comparing Dirichlet normal surface energy of tooth crowns, a new technique of molar shape quantification for dietary inference, with previous methods in isolation and in combination. Am J Phys Anthropol, 145(2), 247–261. https://doi.org/10.1002/ajpa.21489

Burrows, A. M., & Nash, L. T. (2010). Searching for Dental Signals of Exudativory in Galagos. In A. M. Burrows & L. T. Nash (Eds.), The Evolution of Exudativory in Primates (pp. 211– 233). Springer. https://doi.org/10.1007/978-1-4419-6661-2_11

Burrows, A. M., Nash, L. T., Hartstone-Rose, A., Silcox, M. T., López-Torres, S., & Selig, K. R. (2020). Dental Signatures for Exudativory in Living Primates, with Comparisons to Other Gouging Mammals. The Anatomical Record, 303(2), 265–281. https://doi.org/10.1002/ar.24048

Cignoni, P., Callieri, M., Corsini, M., Dellepiane, M., Ganovelli, F., & Ranzuglia, G. (2008). MeshLab: An Open-Source Mesh Processing Tool. Eurographics Italian Chapter Conference.

Cooke, S. B. (2011). Paleodiet of extinct platyrrhines with emphasis on the Caribbean forms: Three-dimensional geometric morphometrics of mandibular second molars. Anat Rec (Hoboken), 294(12), 2073–2091. https://doi.org/10.1002/ar.21502

Datzmann, T., von Helversen, O., & Mayer, F. (2010). Evolution of nectarivory in phyllostomid bats (Phyllostomidae Gray, 1825, Chiroptera: Mammalia). BMC Evol Biol, 10(165).

DeCasien, A. R., Williams, S. A., & Higham, J. P. (2017). Primate brain size is predicted by diet but not sociality. Nature Ecology & Evolution, 1(5), 0112. https://doi.org/10.1038/s41559-017-0112

Dumont, E. R. (1995). Enamel Thickness and Dietary Adaptation among Extant Primates and Chiropterans. Journal of Mammalogy, 76(4), 1127–1136. https://doi.org/10.2307/1382604

Dumont, E. R. (2007). Feeding mechanisms in bats: Variation within the constraints of flight. Integrative and Comparative Biology, 47(1), 137–146. https://doi.org/10.1093/icb/icm007

Dumont, E. R., Davalos, L. M., Goldberg, A., Santana, S. E., Rex, K., & Voigt, C. C. (2012). Morphological innovation, diversification and invasion of a new adaptive zone. Proc Biol Sci, 279(1734), 1797–1805. https://doi.org/10.1098/rspb.2011.2005

Duque-Márquez, A., Ruiz-Ramoni, D., Ramoni-Perazzi, P., & Muñoz-Romo, M. (2019). Bat Folivory in Numbers: How Many, How Much, and How Long? Acta Chiropterologica. https://agris.fao.org/agris-search/search.do?recordID=US202000065528

Evans, A. R., Wilson, G. P., Fortelius, M., & Jernvall, J. (2007). High-level similarity of dentitions in carnivorans and rodents. Nature, 445(7123), 78–81. https://doi.org/10.1038/nature05433

Fedorov, A., Beichel, R., Kalpathy-Cramer, J., Finet, J., Fillion-Robin, J. C., Pujol, S., Bauer, C., Jennings, D., Fennessy, F., Sonka, M., Buatti, J., Aylward, S., Miller, J. V., Pieper, S., & Kikinis, R. (2012). 3D Slicer as an image computing platform for the Quantitative Imaging Network. Magn Reson Imaging, 30(9), 1323–1341. https://doi.org/10.1016/j.mri.2012.05.001

Freeman, P. (1981). Correspondence of Food Habits and Morphology in Insectivorous Bats. Mammalogy Papers: University of Nebraska State Museum. https://digitalcommons.unl.edu/museummammalogy/17

Freeman, P. (1988). Frugivorous and animalivorous bats (Microchiroptera): Dental and cranial adaptations. Mammalogy Papers: University of Nebraska State Museum. https://digitalcommons.unl.edu/museummammalogy/21

Freeman, P. W. (1995). Nectarivorous feeding mechanisms in bats. Biological Journal of the Linnean Society, 56(3), 439–463. https://doi.org/10.1006/bijl.1995.0080

Fulwood, E. L., Shan, S., Winchester, J. M., Gao, T., Kirveslahti, H., Daubechies, I., & Boyer, D. M. (undefined/ed). Reconstructing dietary ecology of extinct strepsirrhines (Primates, Mammalia) with new approaches for characterizing and analyzing tooth shape. Paleobiology, 1–20. https://doi.org/10.1017/pab.2021.9

Gaudioso, P. J., Martínez, J. J., Barquez, R. M., & Díaz, M. M. (2020). Evolution of scapula shape in several families of bats (Chiroptera, Mammalia). Journal of Zoological Systematics and Evolutionary Research, n/a(n/a). https://doi.org/10.1111/jzs.12383

Giannini, N. P., Amador, L. I., & Moyers Arévalo, R. L. (2020). The Evolution of Body Size in Noctilionoid Bats. In T. H. Fleming, L. M. Dávalos, & M. A. R. Mello (Eds.), Phyllostomid Bats: A Unique Mammalian Radiation (p. 123).

Guillerme, T. (2018). dispRity: A modular R package for measuring disparity. Methods in Ecology and Evolution, 9(7), 1755–1763. https://doi.org/10.1111/2041-210X.13022

Hall, R. P., Mutumi, G. L., Hedrick, B. P., Yohe, L. R., Sadier, A., Davies, K. T. J., Rossiter, S. J., Sears, K., Dávalos, L. M., & Dumont, E. R. (n.d.). Find the Food First: An Omnivorous Sensory Morphotype Predates Biomechanical Specialization for Plant Based Diets in Phyllostomid Bats. Evolution, n/a(n/a). https://doi.org/10.1111/evo.14270

Lazzari, V., Charles, C., Tafforeau, P., Vianey-Liaud, M., Aguilar, J.-P., Jaeger, J.-J., Michaux, J., & Viriot, L. (2008). Mosaic Convergence of Rodent Dentitions. PLoS ONE, 3(10), e3607. https://doi.org/10.1371/journal.pone.0003607

Ledogar, J. A., Luk, T. H. Y., Perry, J. M. G., Neaux, D., & Wroe, S. (2018). Biting mechanics and niche separation in a specialized clade of primate seed predators. PLoS ONE, 13(1), e0190689. https://doi.org/10.1371/journal.pone.0190689

Ledogar, J. A., Winchester, J. M., St. Clair, E. M., & Boyer, D. M. (2013). Diet and dental topography in pitheciine seed predators. American Journal of Physical Anthropology, 150(1), 107–121. https://doi.org/10.1002/ajpa.22181

Leiser-Miller, L. B., & Santana, S. E. (2020). Morphological Diversity in the Sensory System of Phyllostomid Bats: Implications for Acoustic and Dietary Ecology. Functional Ecology, n/a(n/a). https://doi.org/10.1111/1365-2435.13561

López-Aguirre, C., Czaplewski, N. J., Link, A., Takai, M., & Hand, S. J. (2020). Dietary and body mass reconstruction of the Miocene neotropical bat Notonycteris magdalenensis (Phyllostomidae) from La Venta, Colombia. BioRxiv, 2020.12.09.418491. https://doi.org/10.1101/2020.12.09.418491

López-Aguirre, C., Hand, S. J., Koyabu, D., Tu, V. T., & Wilson, L. A. B. (2021). Phylogeny and foraging behaviour shape modular morphological variation in bat humeri. Journal of Anatomy, 238(6), 1312–1329. https://doi.org/10.1111/joa.13380

López-Torres, S., Selig, K. R., Prufrock, K. A., Lin, D., & Silcox, M. T. (2018). Dental topographic analysis of paromomyid (Plesiadapiformes, Primates) cheek teeth: More than 15 million years of changing surfaces and shifting ecologies. Historical Biology, 30(1–2), 76–88. https://doi.org/10.1080/08912963.2017.1289378

Makedonska, J., Wright, B. W., & Strait, D. S. (2012). The effect of dietary adaption on cranial morphological integration in capuchins (order Primates, genus Cebus). PLoS ONE, 7(10), e40398. https://doi.org/10.1371/journal.pone.0040398

Melstrom, K. M., & Irmis, R. B. (2019). Repeated Evolution of Herbivorous Crocodyliforms during the Age of Dinosaurs. Current Biology, 29(14), 2389–2395.e3. https://doi.org/10.1016/j.cub.2019.05.076

Monteiro, L. R., & Nogueira, M. R. (2011). Evolutionary patterns and processes in the radiation of phyllostomid bats. BMC Evol Biol, 11, 137. https://doi.org/10.1186/1471-2148-11-137

Morales, A. E., Ruedi, M., Field, K., & Carstens, B. C. (2019). Diversification rates have no effect on the convergent evolution of foraging strategies in the most speciose genus of bats, Myotis. Evolution, 73(11), 2263–2280. https://doi.org/10.1111/evo.13849

Moyers Arévalo, R. L., Amador, L. I., Almeida, F. C., & Giannini, N. P. (2018). Evolution of Body Mass in Bats: Insights from a Large Supermatrix Phylogeny. Journal of Mammalian Evolution. https://doi.org/10.1007/s10914-018-9447-8

Nogueira, M. R., & Peracchi, A. L. (2003). Fig-Seed Predation by 2 Species of Chiroderma: Discovery of a New Feeding Strategy in Bats. Journal of Mammalogy, 84(1), 225–233. https://doi.org/10.1644/1545-1542(2003)084<0225:FSPBSO>2.0.CO;2

Nogueira, M. R., Peracchi, A. L., & Monteiro, L. R. (2009). Morphological correlates of bite force and diet in the skull and mandible of phyllostomid bats. Functional Ecology, 23(4), 715– 723. https://doi.org/10.1111/j.1365-2435.2009.01549.x

Pampush, J. D., Spradley, J. P., Morse, P. E., Harrington, A. R., Allen, K. L., Boyer, D. M., & Kay, R. F. (2016). Wear and its effects on dental topography measures in howling monkeys (Alouatta palliata). American Journal of Physical Anthropology, 161(4), 705–721. https://doi.org/10.1002/ajpa.23077

Pampush, J. D., Winchester, J. M., Morse, P. E., Vining, A. Q., Boyer, D. M., & Kay, R. F. (2016). Introducing molaR: a New R Package for Quantitative Topographic Analysis of Teeth (and Other Topographic Surfaces). Journal of Mammalian Evolution, 23(4), 397–412. https://doi.org/10.1007/s10914-016-9326-0

Pérez-Ramos, A., Romero, A., Rodriguez, E., & Figueirido, B. (2020). Three-dimensional dental topography and feeding ecology in the extinct cave bear. Biology Letters. https://doi.org/10.1098/rsbl.2020.0792

Pineda-Munoz, S., Lazagabaster, I. A., Alroy, J., & Evans, A. R. (2017). Inferring diet from dental morphology in terrestrial mammals. Methods in Ecology and Evolution, 8(4), 481–491. https://doi.org/10.1111/2041-210X.12691

Prufrock, K. A., Boyer, D. M., & Silcox, M. T. (2016). The first major primate extinction: An evaluation of paleoecological dynamics of North American stem primates using a homology free measure of tooth shape. Am J Phys Anthropol, 159(4), 683–697. https://doi.org/10.1002/ajpa.22927

Revell, L. J. (2012). phytools: An R package for phylogenetic comparative biology (and other things). Methods in Ecology and Evolution, 3(2), 217–223. https://doi.org/10.1111/j.2041-210X.2011.00169.x

Rojas, D., Pereira, M. J. R., Fonseca, C., & Dávalos, L. M. (2018). Eating down the food chain: Generalism is not an evolutionary dead end for herbivores. Ecology Letters, 21(3), 402– 410. https://doi.org/10.1111/ele.12911

Rojas, D., Vale, Á., Ferrero, V., & Navarro, L. (2011). When did plants become important to leaf-nosed bats? Diversification of feeding habits in the family Phyllostomidae. Molecular Ecology, 20(10), 2217–2228. https://doi.org/10.1111/j.1365-294X.2011.05082.x

Rojas, D., Vale, Á., Ferrero, V., & Navarro, L. (2012). The role of frugivory in the diversification of bats in the Neotropics. Journal of Biogeography, 39(11), 1948–1960. https://doi.org/10.1111/j.1365-2699.2012.02709.x

Rojas, D., Warsi, O. M., & Dávalos, L. M. (2016). Bats (Chiroptera: Noctilionoidea) Challenge a Recent Origin of Extant Neotropical Diversity. Systematic Biology, 65(3), 432–448. https://doi.org/10.1093/sysbio/syw011

Rossoni, D. M., Assis, A. P. A., Giannini, N. P., & Marroig, G. (2017). Intense natural selection preceded the invasion of new adaptive zones during the radiation of New World leaf-nosed bats. Scientific Reports, 7. https://doi.org/ARTN 11076 10.1038/s41598-017-08989-6

Rossoni, D. M., Costa, B. M. A., Giannini, N. P., & Marroig, G. (2019). A multiple peak adaptive landscape based on feeding strategies and roosting ecology shaped the evolution of cranial covariance structure and morphological differentiation in phyllostomid bats. Evolution, 73(5), 961–981. https://doi.org/10.1111/evo.13715

Santana, S. E., & Cheung, E. (2016). Go big or go fish: Morphological specializations in carnivorous bats. Proceedings of the Royal Society B: Biological Sciences, 283(1830), 20160615. https://doi.org/10.1098/rspb.2016.0615

Santana, S. E., & Dumont, E. R. (2009). Connecting behaviour and performance: The evolution of biting behaviour and bite performance in bats. Journal of Evolutionary Biology, 22(11), 2131–2145. https://doi.org/10.1111/j.1420-9101.2009.01827.x

Santana, S. E., Grosse, I. R., & Dumont, E. R. (2012). Dietary hardness, loading behavior, and the evolution of skull form in bats. Evolution, 66(8), 2587–2598. https://doi.org/10.1111/j.1558-5646.2012.01615.x

Santana, S. E., Strait, S., & Dumont, E. R. (2011). The better to eat you with: Functional correlates of tooth structure in bats. Functional Ecology, 25, 839–847. https://doi.org/10.1111/j.1365-2435.2011.01832.x

Selig, K. R., Khalid, W., & Silcox, M. T. (2021). Mammalian molar complexity follows simple, predictable patterns. Proceedings of the National Academy of Sciences, 118(1). https://doi.org/10.1073/pnas.2008850118

Selig, K. R., López-Torres, S., Sargis, E. J., & Silcox, M. T. (2019). First 3D Dental Topographic Analysis of the Enamel-Dentine Junction in Non-Primate Euarchontans: Contribution of the Enamel-Dentine Junction to Molar Morphology. Journal of Mammalian Evolution, 26(4), 587–598. https://doi.org/10.1007/s10914-018-9440-2

Selig, K. R., Sargis, E. J., Chester, S. G. B., & Silcox, M. T. (2020). Using three-dimensional geometric morphometric and dental topographic analyses to infer the systematics and paleoecology of fossil treeshrews (Mammalia, Scandentia). Journal of Paleontology, 94(6), 1202–1212. https://doi.org/10.1017/jpa.2020.36

Shi, J. J., & Rabosky, D. L. (2015). Speciation dynamics during the global radiation of extant bats. Evolution, 69(6), 1528–1545. https://doi.org/10.1111/evo.12681

Shi, J. J., Westeen, E. P., & Rabosky, D. L. (2018). Digitizing extant bat diversity: An open-access repository of 3D μCT-scanned skulls for research and education. PLoS ONE, 13(9), e0203022. https://doi.org/10.1371/journal.pone.0203022

Ungar, P. (2004). Dental topography and diets of Australopithecus afarensis and early Homo. J Hum Evol, 46(5), 605–622. https://doi.org/10.1016/j.jhevol.2004.03.004

Ungar, P. S., Healy, C., Karme, A., Teaford, M., & Fortelius, M. (2018). Dental topography and diets of platyrrhine primates. Historical Biology, 30(1–2), 64–75. https://doi.org/10.1080/08912963.2016.1255737

Ungar, P. S., & M’Kirera, F. (2003). A solution to the worn tooth conundrum in primate functional anatomy. Proceedings of the National Academy of Sciences, 100(7), 3874– 3877. https://doi.org/10.1073/pnas.0637016100

Villalobos-Chaves, D., Padilla-Alvárez, S., & Rodríguez-Herrera, B. (2016). Seed predation by the wrinkle-faced bat Centurio senex: A new case of this unusual feeding strategy in Chiroptera. Journal of Mammalogy, 97(3), 726–733. https://doi.org/10.1093/jmammal/gyv222

Winchester, J. M., Boyer, D. M., St Clair, E. M., Gosselin-Ildari, A. D., Cooke, S. B., & Ledogar, J. A. (2014). Dental topography of platyrrhines and prosimians: Convergence and contrasts. Am J Phys Anthropol, 153(1), 29–44. https://doi.org/10.1002/ajpa.22398

Zuercher, M. E., Monson, T. A., Dvoretzky, R. R., Ravindramurthy, S., & Hlusko, L. J. (2020). Dental Variation in Megabats (Chiroptera: Pteropodidae): Tooth Metrics Correlate with Body Size and Tooth Proportions Reflect Phylogeny. Journal of Mammalian Evolution. https://doi.org/10.1007/s10914-020-09508-7

